# Gut delivery of pentameric GLP-1 using genetically engineered *Bacillus subtilis*

**DOI:** 10.1101/2025.10.07.680898

**Authors:** Ningyuan Ye, Francesco Di Pierro, Nicola Zerbinati, Maria Laura Tanda, Chenglong Duan, Jingchen Xu, Jure Zupet, Jiahe Li

**Affiliations:** Department of Biomedical Engineering, College of Engineering and School of Medicine, University of Michigan, Michigan, USA; Microbiota International Clinical Society, Turin, Italy; Scientific & Research Department, Velleja Research, Milan, Italy; Endocrine Unit, Department of Medicine and Surgery, University of Insubria, Varese, Italy; Department of Medicine and Technological Innovation, University of Insubria, Varese, Italy; Qingdao Saiding Biological Pharmaceutical Co., Shandong Province, China; Research and Development, Eustone, Slovenia; Department of Biochemistry, University of Michigan, Michigan, USA

**Keywords:** Glucagon-like Peptide-1, *Bacillus subtilis*, Synthetic Biology, Genetically-modified microorganisms, Gut Delivery

## Abstract

Glucagon-like peptide-1 (GLP-1) receptor agonists and GLP-1 analogs are widely used for type 2 diabetes and weight management, but current approaches can be limited by manufacturing complexity, cost, formulation requirements, and the need for repeated administration. Living microbial delivery systems offer a potential strategy for sustained gastrointestinal production of therapeutic peptides, yet achieving stable expression, intestinal survival, and biologically relevant systemic exposure remains challenging. Here, we developed an engineered *Bacillus subtilis* (*B. subtilis*) spore-based platform for gastrointestinal delivery of pentameric GLP-1. By leveraging the stress resistance, storage stability, and genetic tractability of *B. subtilis*, we generated a chromosomally integrated strain that secretes a multimeric GLP-1 construct, which is processed in the intestine to release active GLP-1 monomers. Oral administration of engineered spores resulted in detectable serum GLP-1-related peptide signals in mice, accompanied by preliminary glucose-lowering activity in a fasting-refeeding test. In a longer-term study, engineered spore administration was associated with reduced body weight and adiposity in diet-induced obese mice, with semaglutide included as a positive pharmacologic benchmark. To improve translational suitability, we further generated a marker-free strain, which retained probiotic-relevant features of the parental strain, including gastrointestinal stress tolerance, epithelial cell adhesion, and antibiotic susceptibility. These findings establish proof-of-concept evidence that engineered spore-forming bacteria may serve as a scalable gastrointestinal peptide-delivery platform and support further evaluation of this approach in disease-relevant models of metabolic dysfunction.

## Introduction

Glucagon-like peptide-1 (GLP-1) is an endogenous incretin hormone secreted primarily by intestinal L cells and plays a central role in metabolic regulation.^[1–3]^ Biologically active GLP-1 stimulates glucose-dependent insulin secretion, delays gastric emptying, suppresses appetite, and contributes to body-weight control and lipid metabolism.^[4–6]^ These pleiotropic activities have made the GLP-1 pathway a major therapeutic target, and GLP-1 receptor agonists and GLP-1 analogs are now widely used for type 2 diabetes and weight management.^[7, 8]^ Beyond glycemic control, GLP-1-based therapies are also being investigated for broader metabolic indications, including obesity-associated metabolic dysfunction and non-alcoholic fatty liver disease.^[9–11]^

Despite their clinical impact, GLP-1-based therapies face important delivery and manufacturing challenges. Native GLP-1 is rapidly degraded by dipeptidyl peptidase-4 (DPP-4), with a circulating half-life of only approximately 1–2 minutes, limiting its direct therapeutic utility.^[12–14]^ To overcome this limitation, clinically used GLP-1 receptor agonists and analogs incorporate peptide engineering, amino acid substitutions, lipidation, albumin binding, PEGylation, or other chemical modifications to prolong stability and systemic exposure. Additional delivery strategies, including absorption enhancers, liposomal formulations, and polymer-based nanoparticles, have also been explored to protect GLP-1 peptides from gastrointestinal degradation and improve bioavailability.^[15–19]^ Although these approaches have substantially advanced the field, they can involve complex manufacturing processes, high production costs, specialized formulation requirements, and repeated administration.

Engineered microbial systems offer a potential alternative strategy for gastrointestinal production and delivery of therapeutic peptides. In principle, living microbial delivery platforms could provide scalable production, local synthesis within the gastrointestinal tract, and sustained exposure to biologically active molecules.^[20, 21]^ However, successful development of such systems requires overcoming several barriers, including stable genetic expression, survival during gastrointestinal transit, efficient secretion of the therapeutic product, and achievement of biologically relevant systemic exposure.^[22]^ These challenges are particularly important for peptide hormones such as GLP-1, which are susceptible to proteolytic degradation and require appropriate processing to generate active forms.

A previous microbial delivery strategy used recombinant *Lactobacillus* for oral delivery of a pentameric GLP-1 construct in diabetic rats. In this design, GLP-1 monomers were arranged in a pentameric form that could be cleaved by intestinal trypsin to release active GLP-1 peptides capable of crossing the intestinal mucosa and entering the bloodstream.^[23]^ Although purified pentameric GLP-1 stimulated insulin secretion in vitro, in vivo administration of recombinant *Lactobacillus* produced limited efficacy, likely due to plasmid instability in the absence of selective pressure, reduced bacterial viability during gastrointestinal transit, and insufficient in situ production of the recombinant product. These limitations highlight the need for microbial chassis with improved gastrointestinal robustness and genetic stability.

To address these challenges, we adapted the pentameric GLP-1 delivery concept to *Bacillus subtilis* (*B. subtilis*), a genetically tractable, Gram-positive bacterium commonly found in nature, fermented foods, and probiotic formulations. *B. subtilis* is generally regarded as safe and can form stress-resistant spores with strong tolerance to heat, storage, transport, acidic conditions, and other stresses encountered during passage through the upper gastrointestinal tract. These properties make *B. subtilis* an attractive chassis for oral microbial delivery.^[24, 25]^ In addition, chromosomal integration of expression cassettes can support stable recombinant protein production while reducing the risk of plasmid loss in the absence of selective pressure.^[26]^

Here, we developed an engineered *B. subtilis* spore-based platform for gastrointestinal delivery of pentameric GLP-1. We designed a chromosomally integrated expression cassette encoding a secreted pentameric GLP-1 construct intended for intestinal processing into active GLP-1 monomers. We evaluated whether oral administration of engineered spores could increase serum GLP-1-related peptide levels and elicit preliminary metabolic effects in mice. We further assessed longer-term effects on body weight and adiposity in diet-induced obese mice, using semaglutide as a positive pharmacologic benchmark for GLP-1 pathway activation rather than as a direct delivery-platform comparator. Finally, to improve translational suitability, we generated a marker-free strain and evaluated its probiotic-relevant properties, including gastrointestinal stress tolerance, epithelial adhesion, and antibiotic susceptibility. Together, this study establishes a proof-of-concept microbial platform for gastrointestinal GLP-1 delivery and supports further evaluation of engineered spore-forming bacteria in disease-relevant models of metabolic dysfunction, without implying that the present data demonstrates treatment of diabetes.

## Results

### Engineering a spore-forming bacterial platform for GLP-1 delivery

We selected the genetically tractable *B. subtilis* PY79 strain as a chassis for developing a gastrointestinal GLP-1 delivery platform because of its non-pathogenic profile, ability to form stress-resistant spores, and suitability for stable chromosomal engineering. To support robust production of the recombinant peptide, we used the strong promoter PNBP3510 to drive expression from a single chromosomal copy. The GLP-1 construct was fused to the AmyQ signal peptide to enable Sec-dependent secretion into the extracellular environment. Because direct expression of monomeric GLP-1 was insufficient in preliminary expression studies, we designed a pentameric GLP-1 construct in which five GLP-1 units were linked in tandem. To further optimize intestinal trypsin-mediated processing, we generated two pentameric GLP-1 variants. The wild-type pentamer retained the native GLP-1 sequence and consisted of five directly linked GLP-1 monomers arranged in tandem. In contrast, the intestinal trypsin-resistant (Tr) pentameric GLP-1 (5×GLP-1) construct contained engineered amino acid substitutions (A2G, K20Q, and K28D) at internal trypsin-cleavage sites within individual GLP-1 monomers, thereby restricting trypsin-mediated cleavage to the junctions between adjacent GLP-1 units (**Figure 1A**). As a result, the wild-type construct served as a control that was susceptible to internal degradation, whereas the Tr construct preferentially released intact GLP-1 monomers following trypsin digestion (**Figure 1B**). The expression cassette contained the PNBP3510 promoter, the AmyQ secretion signal, and either wild-type or Tr pentameric GLP-1 sequences, with or without a C-terminal FLAG tag, and all DNA fragments were generated by polymerase chain reaction (PCR) (**Figure 1C**). The cassette was assembled with homology arms targeting the non-essential *amyE* locus and integrated into the *B. subtilis* PY79 chromosome by homologous recombination. Expression of the GLP-1 constructs was validated by sodium dodecyl sulfate-polyacrylamide gel electrophoresis (SDS-PAGE) and Western blot analysis (**Figure S1**). The trypsin-resistant pentameric GLP-1 strain, PY79 Tr 5×GLP-1, was selected for downstream in vivo studies. A recombinant PY79 strain expressing green fluorescent protein (GFP) was generated and used as a bacterial control to account for the effects of oral spore administration independent of GLP-1 expression.

**Figure 1.**
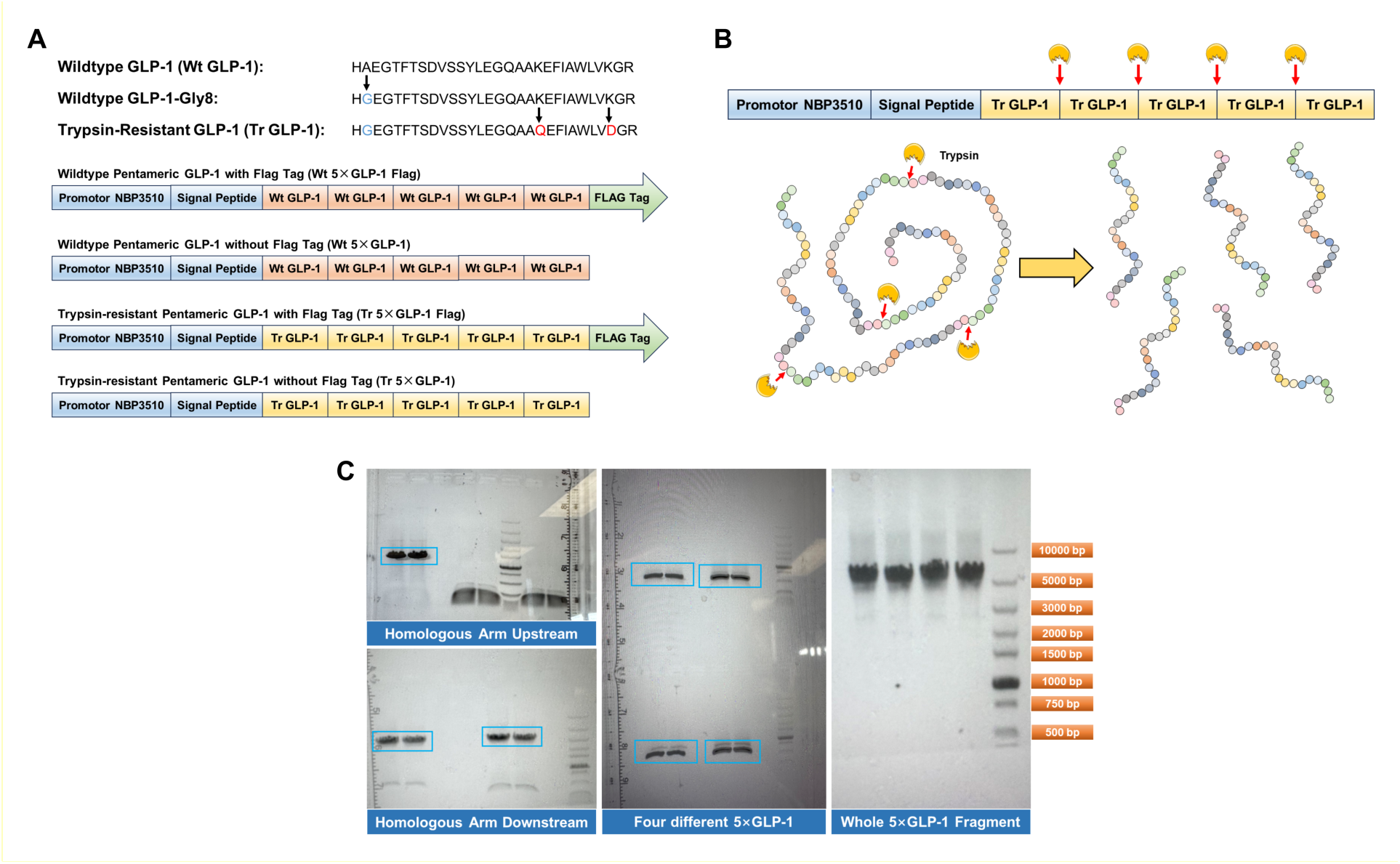
Design and construction of pentameric GLP-1 expression cassettes. (**A**) Schematic representation of wild-type GLP-1 and trypsin-resistant pentameric GLP-1 constructs with or without a C-terminal FLAG tag. The constructs contain the PNBP3510 promoter, AmyQ signal peptide, and five tandem GLP-1 repeats. (**B**) Schematic illustration of trypsin-mediated processing of Tr 5×GLP-1 into 5 intact monomeric GLP-1 peptides. (**C**) PCR amplification and validation of homologous arms and pentameric GLP-1 fragments used for chromosomal integration into the *amyE* locus of *B. subtilis* PY79. Representative PCR gel images from three independent experiments showing consistent results.

### Oral administration of engineered spores enables intestinal recovery, systemic GLP-1 exposure, and shows preliminary glucose-lowering activity

We next evaluated whether orally administered engineered *B. subtilis* spores could be recovered from the gastrointestinal tract and support detectable systemic GLP-1 exposure in mice. Eight-week-old female BALB/c mice were randomized into two groups and administered either PY79 GFP spores or PY79 Tr 5×GLP-1 spores by oral gavage for three consecutive days at a dose of 1 × 10^9^ colony-forming units (CFU) per mouse (**Figure 2A**). Fecal samples collected after treatment were plated on selective medium to confirm recovery of the administered strains from the gastrointestinal tract. Colonies were recovered from fecal samples of both treatment groups, supporting successful passage and recovery of the administered spores following oral dosing (**Figure 2B**).

**Figure 2.**
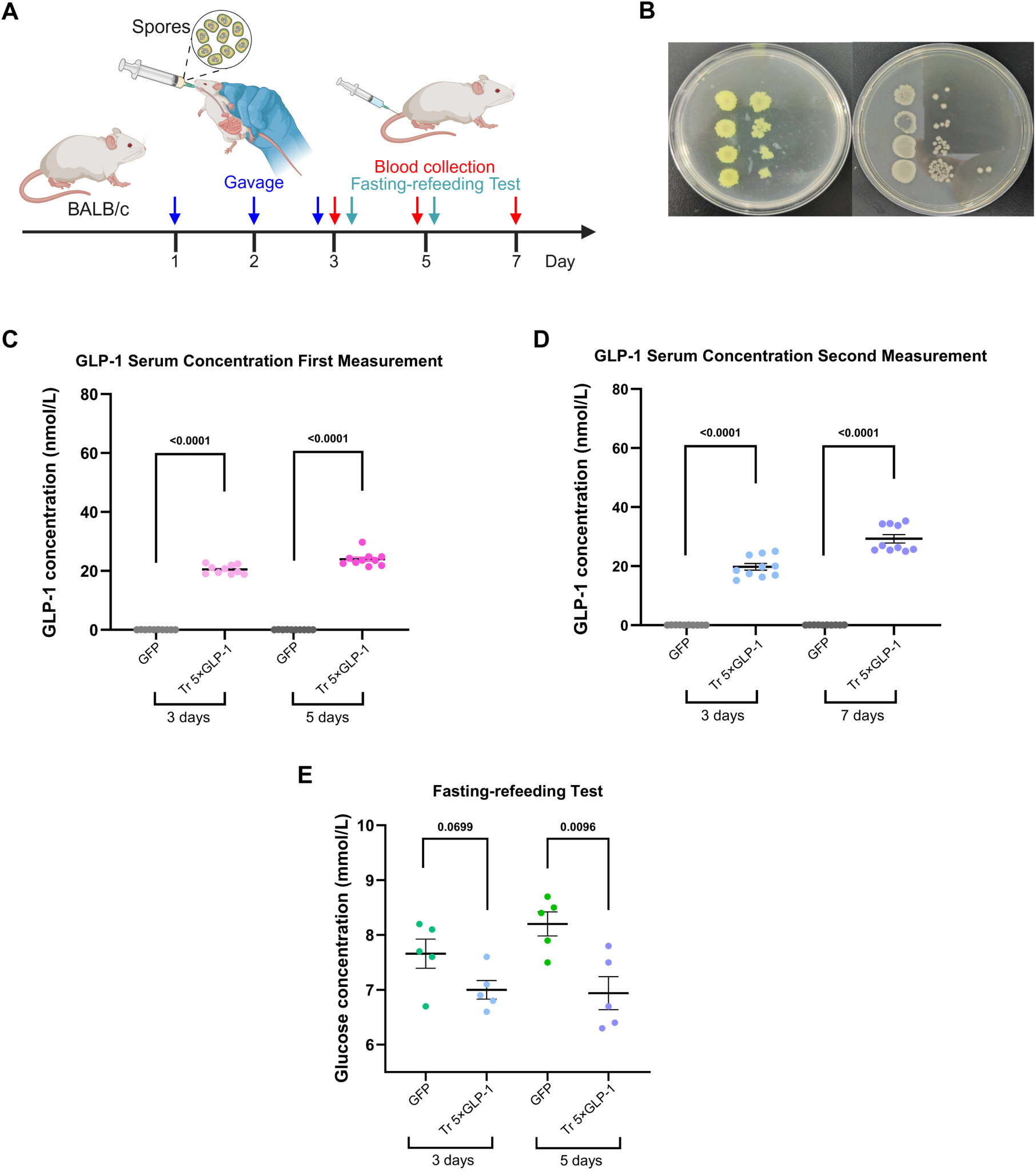
Oral administration of engineered spores enables intestinal recovery, systemic GLP-1 exposure and preliminary glucose-lowering activity. (**A**) Eight-week-old female BALB/c mice received oral gavage of either PY79 GFP spores or PY79 Tr 5×GLP-1 spores for three consecutive days. Blood samples were collected on days 3 and 5 in the first trial and on days 3 and 7 in the second trial. Fasting-refeeding tests were performed on days 3 and 5 in the first trial. (**B**) Representative selective LB agar plates containing chloramphenicol showing fecal bacterial recovery on day 3 from PY79 GFP (left plate) and PY79 Tr 5×GLP-1 (right plate) treatment groups during the first serum GLP-1 measurement trial. (**C**) First serum GLP-1 measurement trial. Blood samples were collected on days 3 and 5 following oral administration. Serum GLP-1-related peptide concentrations were measured in PY79 GFP and PY79 Tr 5×GLP-1 treatment groups. (**D**) Second serum GLP-1 measurement trial. Blood samples were collected on days 3 and 7 following oral administration. Serum GLP-1-related peptide concentrations were measured in PY79 GFP and PY79 Tr 5×GLP-1 treatment groups. (**E**) Fasting-refeeding test. Mice were fasted for 16 h prior to refeeding. Blood glucose levels were measured 2 h after refeeding using tail-vein blood samples and a glucose monitoring kit. Panel **B** shows representative plate images from repeated fecal sampling demonstrating consistent bacterial recovery patterns. Panels **C** and **D** show data from ten biological replicates, and panel **E** shows data from five biological replicates. For **C**, **D** and **E**, data are presented as mean ± SD and statistical analyses were performed using two-way repeated-measures ANOVA followed by Šídák’s multiple-comparisons test comparing GFP and Tr 5×GLP-1 at each timepoint. Panel **A** created with BioRender.com.

Serum GLP-1-related peptide signals were subsequently evaluated in two independent oral administration trials performed using separate cohorts of mice. In the first trial, blood samples were collected on days 3 and 5. Mice receiving PY79 Tr 5×GLP-1 spores showed mean serum GLP-1-related peptide concentrations of 20.61 nM on day 3 and 23.96 nM on day 5, whereas GLP-1 was nearly undetectable in mice receiving PY79 GFP control spores (**Figure 2C**). In the second trial, blood samples were collected on days 3 and 7. Mice treated with PY79 Tr 5×GLP-1 spores showed mean serum GLP-1-related peptide concentrations of 18.72 nM on day 3 and 29.27 nM on day 7, again with minimal detectable GLP-1 in the GFP control group (**Figure 2D**). These findings indicate that oral administration of the engineered GLP-1-producing spores resulted in detectable serum signals for GLP-1-related peptides. Although the exact contribution of intestinal processing, mucosal transport, and systemic stability requires further mechanistic investigation, the marked difference between the Tr 5×GLP-1 and GFP control groups supports GLP-1 exposure associated with the engineered expression cassette rather than spore administration alone.

To determine whether the increase in serum GLP-1 was associated with a measurable physiological effect, we performed a fasting-refeeding test using five mice randomly selected from the first serum GLP-1 measurement trial cohort. Mice were fasted overnight and then refed, and blood glucose levels were measured two hours after food reintroduction. On day 3, mice receiving PY79 Tr 5×GLP-1 spores showed a trend toward lower blood glucose compared with GFP controls, although the difference was not statistically significant. On day 5, mice receiving PY79 Tr 5×GLP-1 spores showed significantly lower blood glucose levels two hours after refeeding compared with mice receiving GFP control spores (**Figure 2E**). Taken together, these data provide preliminary evidence that oral delivery of GLP-1-producing spores can result in glucose-lowering activity following short-term administration.

### Chronic oral administration is associated with reduced body weight and adiposity in diet-induced obese mice

Having observed systemic GLP-1 exposure and preliminary glucose-lowering activity after short-term administration, we next evaluated whether prolonged oral dosing was associated with changes in body weight and adiposity under chronic dietary conditions. Male diet-induced obese mice and age-matched lean C57BL/6J mice at 8 weeks of age maintained on the same high-fat diet were assigned to receive water, GFP control spores, Tr 5×GLP-1 spores, or semaglutide. Both spore-treated groups received 2 × 10^9^ CFU per mouse per day. Body weight and body fat percentage were measured weekly throughout the treatment period (**Figure 3A**). Semaglutide was included as a positive pharmacologic benchmark for GLP-1 pathway activation. In diet-induced obese mice, animals receiving Tr 5×GLP-1 spores showed a progressive reduction in body weight over the treatment period. In contrast, water-treated mice continued to gain weight, whereas GFP spore-treated controls exhibited relatively stable body weights throughout the study period. The body-weight trajectory in the Tr 5×GLP-1 spore-treated group was similar to that observed in the semaglutide benchmark group (**Figure 3B**). A similar pattern was observed for body fat percentage, with Tr 5×GLP-1 spore-treated obese mice showing reduced adiposity compared with water-treated and GFP spore-treated controls (**Figure 3C**). In lean mice, all treatment groups continued to gain weight over time, consistent with normal growth during the study period. However, Tr 5×GLP-1 spore-treated lean mice showed a modest attenuation of body-weight gain and body fat accumulation compared with water-treated and GFP spore-treated controls (**Figure 3D,E**). Collectively, these findings suggest that the metabolic effects associated with GLP-1-producing spores were more pronounced in diet-induced obesity than in lean animals.

**Figure 3.**
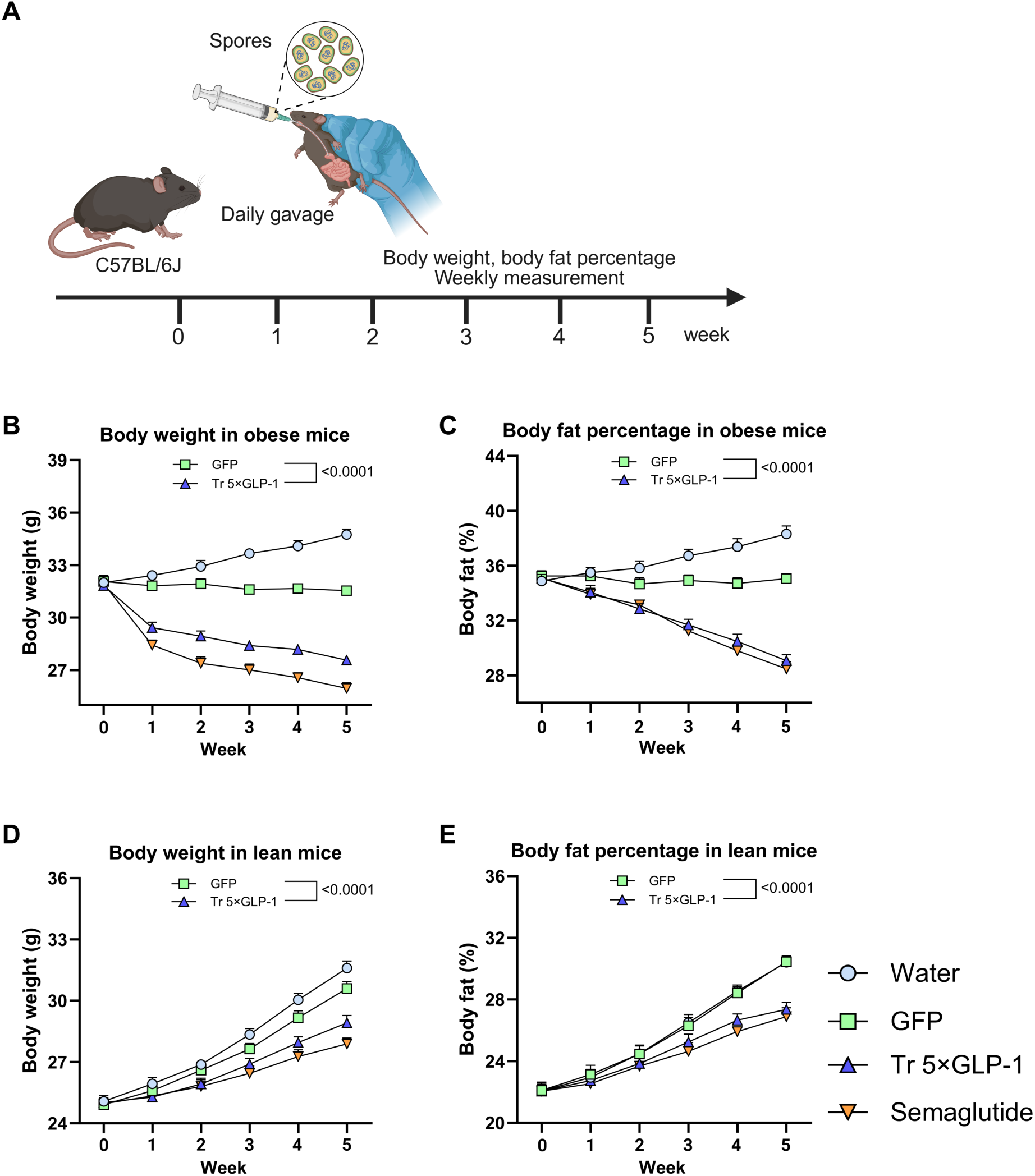
Oral administration of engineered spores is associated with reduced body weight and adiposity. (**A**) Schematic of the long-term oral administration study. Eight-week-old male diet-induced obese and age-matched lean C57BL/6J mice received daily oral gavage of water, GFP control spores, Tr 5×GLP-1 spores, or semaglutide for 5 weeks. Body weight and body fat percentage were measured weekly throughout the treatment period. (**B**) Body weight measurements in obese mice. (**C**) Body fat percentage measurements in obese mice. (**D**) Body weight measurements in lean mice. (**E**) Body fat percentage measurements in lean mice. For panels **B**–**E**, data are presented as mean ± SD from ten biological replicates. Statistical analyses were performed using two-way repeated-measures ANOVA followed by Šídák’s multiple-comparisons test comparing GFP and Tr 5×GLP-1 at each timepoint. Panel **A** created with BioRender.com.

Together, these findings support the potential of engineered *B. subtilis* spores to deliver biologically active GLP-1 in vivo and to influence metabolic readouts after chronic oral administration. However, because the study was not designed as a definitive efficacy study in a diabetes or obesity disease-treatment model, these results should be presented as preliminary metabolic bioactivity supporting further evaluation in disease-relevant models.

### Generation of a marker-free GLP-1-producing strain

To improve the translational suitability of the engineered GLP-1-producing strain, we removed the chloramphenicol resistance cassette located downstream of the 5×GLP-1 expression cassette from PY79 Tr 5×GLP-1 using a Cre/lox recombination strategy. The parental engineered strain contained loxP sites flanking the antibiotic resistance cassette at the chromosomal integration locus. After transformation with the temperature-sensitive Cre-expressing plasmid pDR244 containing spectinomycin resistance cassette, colonies were selected and then cultured under conditions that promoted Cre-mediated recombination and plasmid curing. Candidate colonies that lost both chloramphenicol and spectinomycin resistance were identified by replica plating. PCR and sequencing confirmed successful excision of the chloramphenicol resistance cassette, retention of a single loxP site, and preservation of the adjacent genomic sequence. The resulting marker-free strain was designated JH389 and deposited at the Belgian Coordinated Collections of Microorganisms/Laboratorium voor Microbiologie (BCCM/LMG) collection as *B. subtilis* LMG P-34037. To determine whether antibiotic marker removal affected GLP-1 expression, we compared protein production before and after marker excision. SDS-PAGE analysis showed a band at the expected molecular weight of approximately 15 kDa in GLP-1-expressing samples, indicating that marker removal did not detectably impair secretion of the recombinant pentameric GLP-1 construct (**Figure S2**).

### Marker-free JH389 retains probiotic-relevant and translationally favorable properties

We next evaluated whether genetic engineering and marker removal altered key properties relevant to future development of the engineered strain. First, strain purity and colony morphology were assessed by streaking recovered PY79 and JH389 samples on tryptic soy agar (TSA) plates.

Both strains produced a single expected colony morphology, supporting culture purity prior to downstream characterization (**Figure S3**). This result is better integrated here rather than presented as a separate Results subsection.

We then assessed antibiotic susceptibility profiles for PY79 and JH389. Across the tested antibiotic panel, minimum inhibitory concentration (MIC) values for both strains were generally at or below available microbiological cut-off values for *B. subtilis* as defined by European Food Safety Authority (EFSA) (**Table 1**). The main exception was streptomycin, for which JH389 showed a two-fold increase and PY79 showed a four-fold increase relative to the listed cut-off. Because two-fold variation is commonly observed in MIC testing and is generally considered within expected assay variability, these data support the conclusion that JH389 retained a favorable antibiotic susceptibility profile after marker removal.

**Table 1.** MIC values (expressed as µg/mL) for *B. subtilis* JH389 and *B. subtilis* PY79.

| Antibiotic | *B. subtilis* JH389 | *B. subtilis* PY79 | EFSA Cut-off |
| --- | --- | --- | --- |
| Gentamicin | < 0.5 | < 0.5 | 4 |
| Kanamycin | < 2 | < 2 | 8 |
| Streptomycin | 16 | 32 | 8 |
| Neomycin | < 0.12 | 0.5 | NA |
| Tetracycline | 2 | 2 | 8 |
| Erythromycin | 2 | 0.25 | 4 |
| Clindamycin | 1 | 2 | 4 |
| Chloramphenicol | 2 | 4 | 8 |
| Ampicillin | < 0.03 | < 0.03 | NA |
| Penicillin | < 0.03 | < 0.03 | NA |
| Vancomycin | <0.25 | 0.25 | 4 |
| Quinupristin/Dalfopristin | 1 | 1 | NA |
| Linezolid | 0.5 | 0.5 | NA |
| Trimethoprim | 0.25 | 0.25 | NA |
| Ciprofloxacin | < 0.25 | < 0.25 | NA |
| Rifampicin | 0.25 | 0.25 | NA |
| Amoxicillin/Clavulanate | 0.06 | 0.06 | NA |
| Cefixime | 8 | 8 | NA |

We further evaluated the ability of PY79 and JH389 to tolerate simulated gastrointestinal conditions. In simulated gastric juice (SGJ), both strains showed high survival after 30 and 60 minutes of exposure. PY79 retained 99.4% and 98.4% residual viability at 30 and 60 minutes, respectively, while JH389 retained 99.1% and 97.8% residual viability at the same time points (**Table 2**).

**Table 2.**
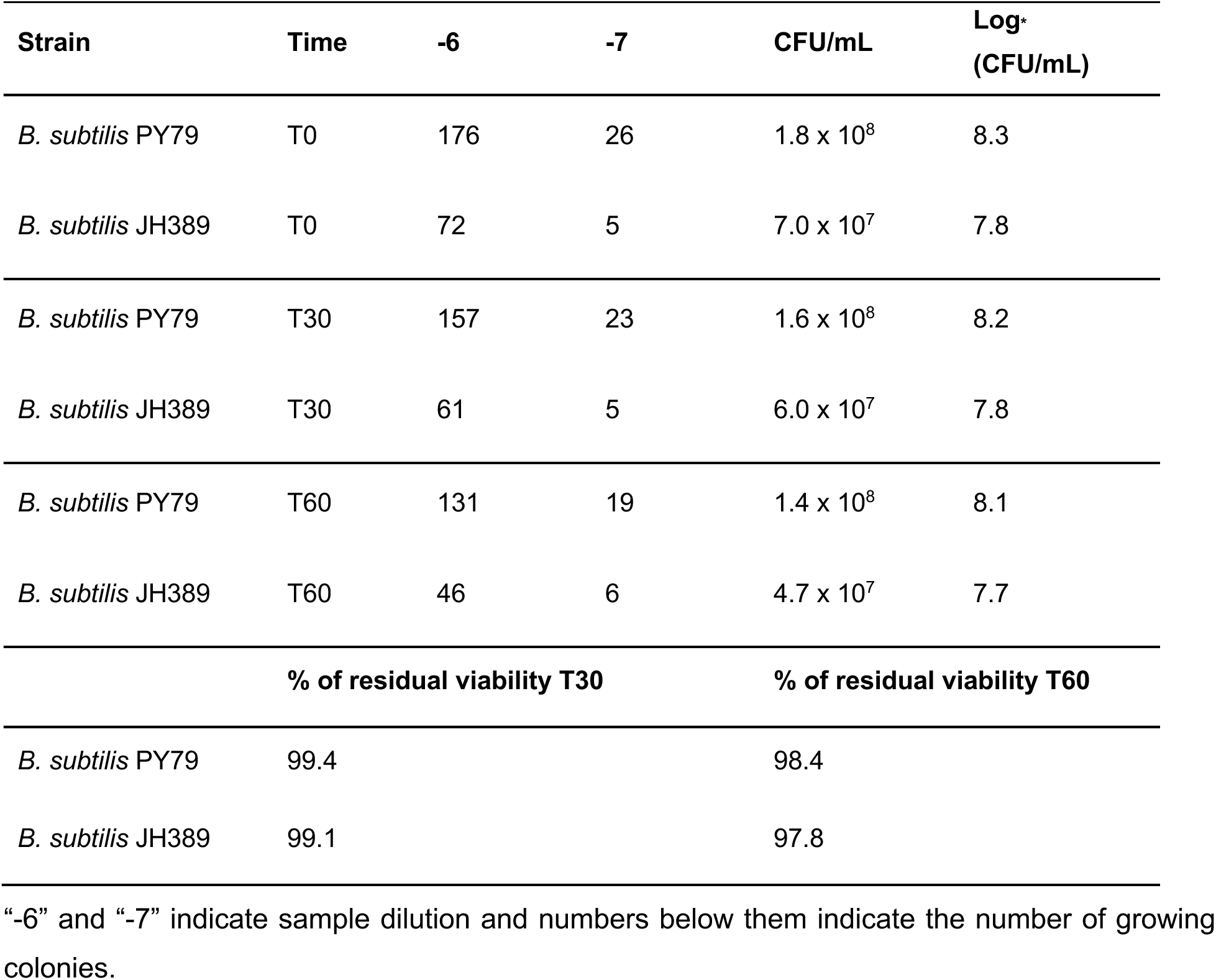
Viable cell counts and log_10_ (CFU/mL) following exposure to SGJ.

In simulated intestinal juice (SIJ) containing pancreatin and bile, both strains also retained substantial viability over extended incubation. PY79 retained 81.2% and 79.1% residual viability at 240 and 360 minutes, respectively, while JH389 retained 84.4% and 81.9% residual viability at the same time points (**Table 3**).

**Table 3.**
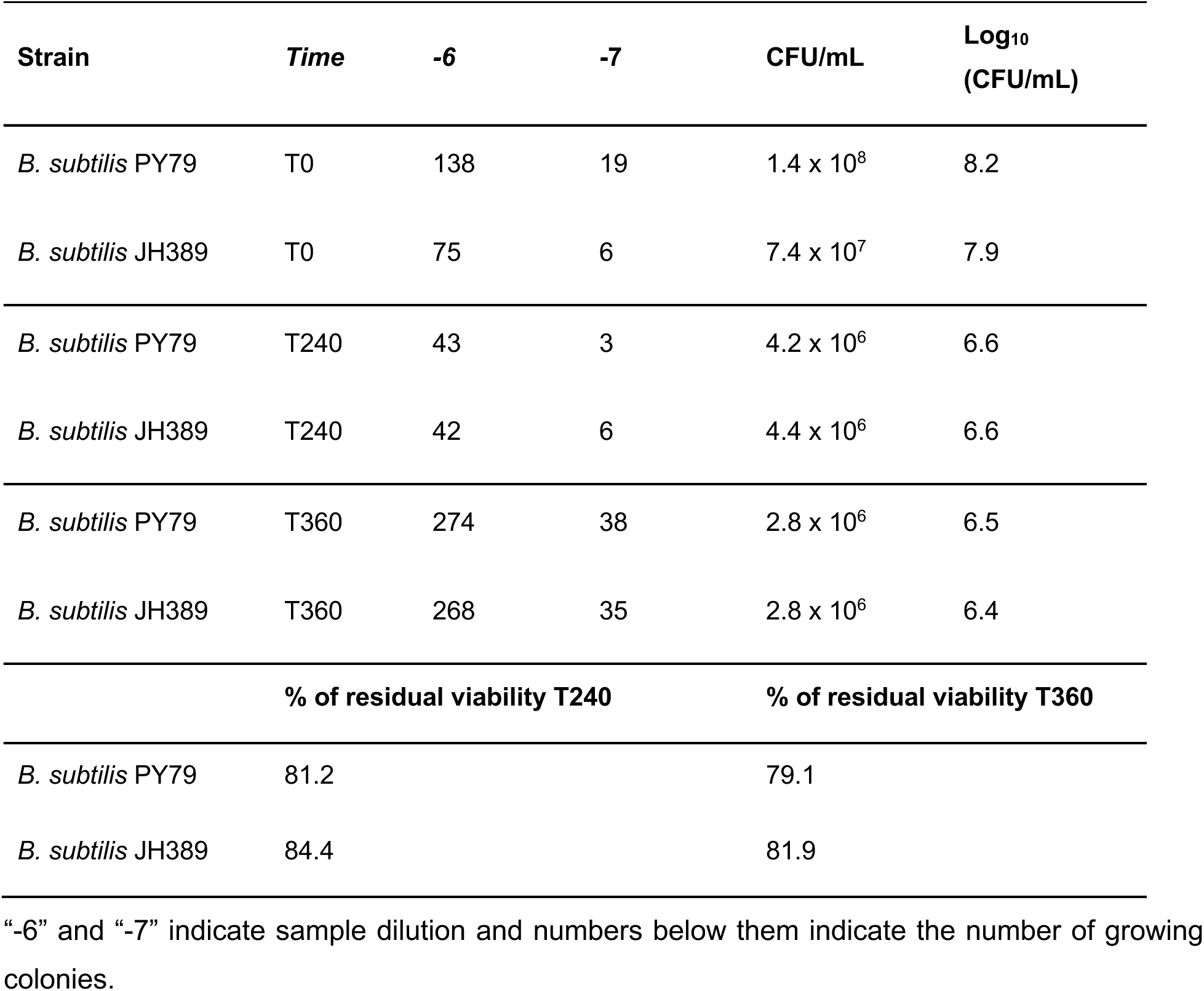
Viable cell counts and log_10_ (CFU/mL) following exposure to SIJ.

Finally, we assessed adhesion to HT29 human intestinal epithelial cells. The technical control strain, *Lacticaseibacillus paracasei* ATCC 334, showed a relative adhesion index of 68%, within the expected assay validation range. PY79 and JH389 also showed measurable adhesion to HT29 cells, with relative adhesion index of 61% and 66%, respectively (**Table 4**).

**Table 4.** Adhesion assay results on HT29 cell line.

| Strain | HT29 cells |  |
| --- | --- | --- |
|  | Inoculated bacteria | Adherent bacteria |

|  | -4 | -5 | CFU/mL | Log<sub>10</sub><br>(CFU) | -2 | -3 | CFU/mL | Log<sub>10</sub><br>(CFU) | Adhesion<br>% |
| --- | --- | --- | --- | --- | --- | --- | --- | --- | --- |
| *L. paracasei*<br>ATCC 334 | 145 | 12 | 1.4 x 10<sup>6</sup> | 6.2 | 163 | 11 | 1.6 x 10<sup>4</sup> | 4.2 | 68 |
| *B. subtilis*<br>PY79 | 108 | 16 | 1.1 x 10<sup>6</sup> | 6.1 | 48 | 7 | 5.0 x 10<sup>3</sup> | 3.7 | 61 |
| *B. subtilis*<br>JH389 | 86 | 6 | 8.4 x 10<sup>5</sup> | 5.9 | 83 | 12 | 8.6 x 10<sup>3</sup> | 3.9 | 66 |

“-4”, “-5”, “-2” and “-3” indicate sample dilution and numbers below them indicate the number of growing colonies.

These values indicate that the marker-free JH389 strain retained epithelial adhesion capacity comparable to the parental PY79 strain. Collectively, these characterization studies show that JH389 retained several probiotic-relevant and translationally favorable features of the parental strain, including expected colony morphology, antibiotic susceptibility, tolerance to simulated gastric and intestinal conditions, and adhesion to intestinal epithelial cells. These results support the use of JH389 as a marker-free GLP-1-producing strain for future studies evaluating engineered spore-forming bacteria as gastrointestinal peptide-delivery platforms.

## Discussion

In this study, we developed a proof-of-concept microbial delivery platform based on engineered *B. subtilis* spores for gastrointestinal production of pentameric GLP-1. By combining chromosomal integration, spore-based oral delivery, and multimeric peptide engineering, we demonstrated that orally administered engineered spores were associated with measurable serum GLP-1-related peptide signals and preliminary metabolic bioactivity in mice. Importantly, the present work should be interpreted as evidence supporting the feasibility of microbial GLP-1 delivery rather than definitive evidence of therapeutic efficacy in diabetes or obesity.

One of the major challenges in microbial peptide delivery is achieving sufficient stability and expression of small bioactive peptides in bacterial systems. Small peptide therapeutics are often susceptible to rapid degradation and inefficient production in prokaryotic hosts, limiting their utility in engineered microbial delivery platforms. To address this limitation, we adopted a pentameric GLP-1 design in which five GLP-1 units were linked in tandem and separated by trypsin-sensitive cleavage sites. This strategy was intended to improve recombinant peptide stability and expression efficiency while permitting intestinal processing into monomeric GLP-1 peptides following oral administration. Our findings suggest that multimeric engineering approaches may represent a useful strategy for microbial production of bioactive peptides that are otherwise difficult to express efficiently in bacterial systems.

Oral administration of engineered spores resulted in detectable serum GLP-1-related peptide signals in both dosing trials, whereas minimal signal was observed in GFP spore-treated controls. Although the precise mechanisms governing intestinal peptide release, epithelial transport, and systemic persistence remain incompletely defined, these data support the feasibility of gastrointestinal peptide production and subsequent systemic exposure following oral administration of engineered spores. Consistent with these observations, the fasting-refeeding test demonstrated preliminary glucose-lowering activity associated with treatment. However, because these studies were performed in non-diabetic mice and were not designed as definitive efficacy studies, the observed metabolic effects should be interpreted cautiously. In longer-term studies, chronic administration of GLP-1-producing spores was associated with sustained reductions in body weight and adiposity in diet-induced obese mice, whereas effects in lean mice were comparatively modest. These findings suggest that metabolic context may influence the magnitude of treatment-associated physiological responses. Collectively, these findings highlight the broader potential of engineered spore-based systems for gastrointestinal peptide delivery and systemic therapeutic biomolecule exposure following oral administration.

To improve translational suitability, we further generated the marker-free strain JH389 through Cre/lox-mediated excision of the chloramphenicol resistance cassette. Removal of antibiotic resistance markers is generally considered favorable for future regulatory development of engineered probiotic systems, as antibiotic resistance genes may facilitate horizontal gene transfer, the dissemination of resistance determinants, or mechanisms associated with extended-spectrum β-lactamases (ESBLs).^[34]^ Importantly, removal of the chloramphenicol resistance cassette did not detectably impair recombinant protein secretion or alter several probiotic-relevant properties of the parental strain, including gastrointestinal stress tolerance, epithelial adhesion, and overall antibiotic susceptibility profiles. These findings support the feasibility of generating genetically engineered *B. subtilis* strains with improved compatibility for future translational applications.

Several limitations of the present study should also be acknowledged. First, the current work primarily establishes proof of concept for microbial GLP-1 delivery and does not constitute a comprehensive therapeutic evaluation in diabetic disease models. Second, the mechanisms governing peptide processing, intestinal absorption, pharmacokinetics, and long-term gastrointestinal persistence remain incompletely characterized. Third, although preliminary metabolic effects were observed, additional studies involving glucose tolerance testing, insulin measurements, food intake analysis, and broader metabolic phenotyping will be required to define the physiological impact of the platform more rigorously. Additionally, because the LC-MS analysis detected GLP-1-related peptide signals under standard serum collection conditions without DPP-4 inhibition, the measured signals may reflect a mixture of engineered peptide fragments, processed active GLP-1, and endogenous host-derived GLP-1 species rather than intact bioactive GLP-1 alone. Additional studies will therefore be required to further define the precise molecular species contributing to the detected serum signals. Future work should also evaluate long-term biosafety, environmental containment strategies, and scalability under translationally relevant manufacturing conditions. Overall, this study establishes engineered spore-forming bacteria as a potential platform for gastrointestinal peptide delivery and supports continued investigation of microbial therapeutics for metabolic and other peptide-based applications.

## Methods and Materials

### Chemicals and reagents

All chemicals and bacterial culture broth were purchased from Fisher Scientific International Inc. (New Hampshire, USA) unless otherwise noted, and were of highest purity or analytical grade.

### Bacterial strains and culture conditions

*B. subtilis* PY79 was grown overnight (30 °C, 220 rpm) in Luria–Bertani (LB) medium, with or without 5 μg/mL chloramphenicol. For downstream applications, overnight bacterial culture was diluted 10 times followed by culturing (37 °C, 220 rpm), until the OD_600_ reached between 0.6 and 0.8. Bacteria were stored in a formulation buffer (2.28 g/L KH_2_PO_4_, 14.5 g/L K_2_HPO_4_, and 15% glycerol, pH 7.5) at 10^11^ CFU/mL, and -80 °C.

### Growth curve measurements

For growth curve analysis, bacterial cultures were initiated in 1 mL of LB medium with or without 5 μg/mL chloramphenicol and incubated overnight at 37 °C with shaking at 220 rpm. Overnight cultures were diluted 1:10, and cell density was measured at OD_600_ using a NanoDrop spectrophotometer. Cultures were then adjusted to an initial OD_600_ of 0.01 in 15 mL of prewarmed, antibiotic-free LB medium in 125 mL flasks. Aliquots (1 mL) were collected at 0–8 h at 1 h intervals for OD_600_ measurement. Growth curves were generated by averaging OD_600_ values from three biological replicates at each time point.

### B. subtilis spore preparation

Overnight bacterial culture was plated on selective 2× SG agar plates (per 1.0 L ddH_2_O; 15.0 g agar, 16.0 g Difco nutrient broth, 0.24 g MgSO_4_, 1.938 g KCl, 19.8 mg MnCl_2_, 0.3 mg FeSO_4_, 0.236 g Ca(NO_3_)_2_, 1.0 g glucose, 143 μL chloramphenicol 35 μg/mL), and incubated at 30 °C for 36 hours. The spore formation rate was examined under a microscope: if the rate exceeded 90%, the spores were scraped and resuspended in ddH_2_O. The spores were then washed by repeated centrifugation (5000 rpm, 10 min, RT) and ultrasonication (1 min sonication 20W, 2100J, 1 min rest, three times per day) for three days. Finally, the spore suspension concentration was quantified via the serial dilution method, and the spores aliquoted at 10^10^ CFU/mL and stored at - 80 °C.

### Homology-directed gene editing

The 31 amino acid sequence of the active form of GLP-1 (7-37) was obtained from the NCBI database. The second amino acid A was mutated to G, and the 20^th^ and 28^th^ amino acids K were mutated to Q and D, respectively. In addition, the last amino acid was removed to obtain a sequence whereby appropriate trypsin cleavage sites would be obtained in the polymeric form of the peptide to preserve the biologically active N-terminal histidine residue on each monomer. This modified sequence was designated as trypsin-resistant GLP-1 (Tr GLP-1), while the unmodified sequence was referred to as wild-type GLP-1 (Wt GLP-1). Each sequence was repeated five times to construct pentameric GLP-1 sequences. The pentameric sequences were designated as Wt 5×GLP-1 for the unmodified sequence, and Tr 5×GLP-1 for the modified trypsin-resistant sequence. Additional variants were designed to include a FLAG tag at the C-terminal end of the pentamer sequence. The sequences were then converted into corresponding nucleotide sequences optimized for *B. subtilis*, and synthesized as gBlock double-stranded DNA fragments (Twist Bioscience Corporation, USA). Similarly, the *amyE* gene sequence was retrieved from NCBI, and primers were designed based on its upstream and downstream homologous regions. The primers were obtained from Integrated DNA Technologies (IDT, USA). DNA sequences for the NBP3510 promoter, the AmyQ signal peptide, terminator regions, and the chloramphenicol resistance marker were readily available in-house. PCR was used to amplify the DNA fragments, which were combined using Gibson assembly.^[27]^

Competent cells of *B. subtilis* PY79 were prepared using 1× MC medium (per 1.0 L ddH_2_O; 10.7 g K_2_HPO_4_, 5.2 g KH_2_PO_4_, 20.0 g glucose, 0.88 g Na_3_C_6_H_5_O_7_·2H_2_O, 0.022 g C_6_H_5_FeNO_7_, 2.2 g C₅H₈KNO₄·H₂O, 1.0 g Casein Hydrolysate, 12.0 g MgSO_4_), and the assembled DNA fragments were transformed into the competent bacteria. The bacteria were then plated on LB media containing chloramphenicol to select potential transformants. The presumed recombinant colonies were restreaked on selective LB plates to propagate the transformants. The resulting colonies were selected and the genetic insert amplified by PCR. The amplicons were further verified by Sanger sequencing (Azenta Life Sciences, USA) and subsequently analysed by alignment to validate the genetic insertions.

### SDS-PAGE and Western blotting

800 μL of recombinant bacterial culture supernatant was subjected to precipitation and concentration using the TCA precipitation method. The precipitated protein samples were resuspended in Laemmli 6× SDS-Sample buffer and denatured by heating at 95 °C for 5 minutes. Two 12% SDS-PAGE gels were prepared for electrophoresis and loaded with protein samples. Following electrophoresis, the first gel was stained with Coomassie Brilliant Blue for one hour, and destained in destaining buffer for one day, while the second gel was used for nitrocellulose membrane transfer. Subsequently, immunoblotting was performed by incubating the nitrocellulose membrane with a Rat-derived anti-FLAG tag antibody (DYKDDDDK, Biolegend, USA) overnight under 4 °C. The membrane was then incubated for 1 h at room temperature with a secondary HRP-conjugated anti-Rat IgG antibody (Cell Signalling Technology, USA). Substrate TMB and hydrogen peroxide were added to generate colorimetric reaction to visualize and confirm the successful expression of the FLAG-tagged genetic construct at the protein level.

### Cre/lox-mediated marker deletion using pDR244

To remove the chloramphenicol resistance cassette flanked by loxP sites at the *amyE* locus, we employed the Cre/lox recombination system.^[28]^ The temperature-sensitive plasmid pDR244 (BGSC, USA), encoding Cre recombinase and conferring spectinomycin resistance, was introduced into the engineered *B. subtilis* strain via transformation. Competent cells were prepared using 1× MC medium, and 500 ng of plasmid DNA was added to 400 μL of competent cells, followed by incubation at 30 °C for 2 hours. Transformants were plated on LB agar supplemented with spectinomycin (50 μg/mL) to select for successful uptake of pDR244. A single spectinomycin-resistant colony was inoculated into antibiotic-free LB broth and incubated at 42 °C overnight to induce Cre expression and simultaneously inhibit plasmid replication, facilitating its curing. The culture was then streaked onto LB agar to isolate single colonies, which were subsequently replica plated onto LB without antibiotic, LB with spectinomycin, and LB with chloramphenicol plates. Colonies that grew only on antibiotic-free LB plates were considered candidate clones for marker excision and plasmid loss. Genomic DNA was extracted from these colonies, and PCR was performed using primers flanking the *amyE* locus and showing in the following sheet (**Table S1**). The PCR product was sequenced to confirm complete deletion of the chloramphenicol cassette and retention of a single loxP site.

### B. subtilis PY79 and B. subtilis JH389 isolation and purity check

A small quantity of freeze-dried powder of each *B. subtilis* PY79 and JH389 strain was streaked onto separate TSA (Oxoid SpA, Rodano, Milan, Italy) plates to isolate the two strains. The plates were incubated under aerobic conditions at 37 °C for 48 hours. After incubation, a visual inspection confirmed the presence of a single colony morphology for both strains, as expected. One representative colony from each *B. subtilis* strain was subsequently picked, re-streaked on fresh TSA plates, and incubated to verify culture purity.

### Antibiotic resistance profiles

Prior to performing the antibiotic susceptibility assay, *B. subtilis* PY79 and *B. subtilis* JH389 strains were propagated on TSA medium and incubated aerobically at 37 °C for 48 hours. Minimal Inhibitory Concentrations (MICs) were determined for eighteen antibiotics: ampicillin, penicillin, clindamycin, linezolid (range: 0.03–16 μg/mL), vancomycin, ciprofloxacin (range: 0.25–128 μg/mL), neomycin, gentamicin, streptomycin (range: 0.5–256 μg/mL), kanamycin (range: 2–1024 μg/mL), erythromycin, quinupristin-dalfopristin (range: 0.016–8 μg/mL), tetracycline, chloramphenicol, rifampicin, and trimethoprim (range: 0.125–64 μg/mL), amoxicillin-clavulanate (range: 0.06–64 μg/mL), and cefixime (range: 0.5–64 μg/mL) using commercially prepared microplates (Sensititre™ EULACBI1 and EULACBI2, ThermoFisher).

Following incubation, strain purity and viability were confirmed. Individual colonies were selected and resuspended in 3 mL of sterile saline solution. Cell suspensions were adjusted to a McFarland standard of 1.0, corresponding to approximately 3.0 × 10^8^ CFU/mL, then diluted 1:1000 in Lactic Streptococcal Medium (LSM) broth. Within 30 minutes from preparation, 100 μL of the diluted bacterial suspension was dispensed into each well of the Sensititre™ microplates using a multichannel pipette. Negative control wells were inoculated with sterile LSM broth only. The Sensititre™ EULACBI1 and EULACBI2 plates contained predefined antibiotic concentrations according to ISO 10932:2010 guidelines.^[29]^ Plates were incubated under aerobic conditions at 37 °C for 48 ± 3 hours. Following incubation, negative control wells were examined for potential contamination. Provided that both positive and negative controls met quality criteria, MIC values were recorded.

### Resistance to simulated gastric juice

The evaluation of resistance to SGJ was conducted in triplicate (if highly comparable, only one test is shown in the Results section) following the internal method IM05, adapted from ISTISAN Reports 2008/36, with minor modifications.^[30]^ *B. subtilis* PY79 and JH389 were first cultured in TSB (Tryptic Soy Broth; Oxoid SpA, Rodano, Milan, Italy) broth. For the assay, cultures were centrifuged, the supernatants discarded, and the resulting pellets resuspended in 5 mL of sterile SGJ (pH 3.4 ± 0.1). SGJ composition included sodium taurocholate (0.08 mM), phospholipids (0.02 mM), sodium (34 mM) and chlorine (59 mM) for a final pH of 3.4 ± 0.1. The suspensions were incubated at 37 °C, and 1 mL aliquots were withdrawn at three timepoints: T0, T30, and T60 minutes. Each sample was serially diluted and plated on TSA to determine total viable counts. Plates were incubated under aerobic conditions at 37 °C for 48 hours. Results are expressed as two percentage values, corresponding to strain survival after 30 and 60 minutes of exposure to SGJ, and are calculated using the formula P = (μ / M) × 100%, where P represents the percentage of resistance of the microbial strain to SGJ; μ is the log_10_-transformed viable cell count after 30 minutes (T30) or 60 minutes (T60) of incubation at 37 °C in SGJ; and M is the log_10_-transformed viable cell count at the initial timepoint (T0), immediately after resuspension in SGJ. This approach enables quantification of each strain’s tolerance to acidic stress over time.

### Resistance to simulated intestinal juice

The evaluation of resistance to SIJ were conducted in triplicate (if highly comparable, only one test is shown in the Results section) using the in-house method IM06 which is accredited in accordance with ISO 17025:2017, with minor modifications.^[30, 31]^ The method is based on viable cell counts performed at defined timepoints: T0 (immediately after resuspension in the SIJ), T240 (after 240 minutes of incubation), and T360 (after 360 minutes of incubation) at 37 °C. *B. subtilis* strains PY79 and JH389 were grown in TSB for 24 hours at 37 °C under aerobic conditions. Cultures were centrifuged, the supernatants discarded, and the bacterial pellets resuspended in 5 mL of SIJ. At each timepoint (T0, T240, and T360), 1 mL of the suspension was collected, serially diluted, and plated on TSA for determination of viable counts. This allowed evaluation of the strains’ survival capacity under simulated intestinal conditions. SIJ composition included pancreatin (1 mM), ox-bile (3 mM) and sodium chloride (9 mM) for a final pH of 8 ± 0.1. All plates were incubated for 48 hours at 37 °C under aerobic conditions. Survival results were expressed as percentage values using the formula P = (μ / M) × 100% where P represents the percentage resistance of the microbial strain to SIJ; μ is the log_10_-transformed viable cell count after 240 minutes (T240) or 360 minutes (T360) of incubation at 37 °C in SIJ; and M is the log_10_-transformed viable cell count at time zero (T0), immediately after resuspension in SIJ. This calculation provides a quantitative estimate of the strain’s resistance to intestinal conditions over time.

### Adhesion to HT29 cell lines

The HT29 human intestinal epithelial cell line was routinely maintained in High Glucose DMEM (Dulbecco’s Modified Eagle Medium) supplemented with 10% heat-inactivated fetal bovine serum (FBS), 50 μg/mL L-glutamine, and 40 μg/mL gentamicin at 37 °C in a 5% CO_2_ atmosphere (all reagents from Euroclone; Pero, Milan, Italy). Two days prior to the adhesion assay, HT29 cells were rinsed with Hank’s Balanced Salt Solution (HBSS), detached using trypsin, counted, and diluted to 2.5 × 10^5^ cells/mL. One well per strain was seeded with the cell suspension in a 24-well plate, which was then incubated at 37 °C with 5% CO_2_ for 48 hours, until confluence was achieved. The day before the assay, *B. subtilis* PY79, *B. subtilis* JH389, and *Lacticaseibacillus paracasei* (*L. paracasei*) ATCC 334 (used as a technical control) were inoculated into TSB (for *B. subtilis*) or de Man, Rogosa and Sharpe (MRS) broth (for *L. paracasei*) and incubated for 24 hours at 37 °C under appropriate atmospheric conditions. On the day of the test, confluence of HT29 monolayers was confirmed. Wells were washed with HBSS and pre-incubated for 1 hour with 875 μL of DMEM High Glucose + 1% FBS at 37 °C, 5% CO_2_. For each sample, an additional well containing only 875 μL of medium (without cells) was included as a control. Meanwhile, the bacterial cultures were centrifuged and washed twice with sterile distilled water. Cell pellets were resuspended and adjusted to McFarland standard 0.5, corresponding to approximately 1.5 × 10^8^ CFU/mL. These suspensions were further diluted 1:10 in High Glucose DMEM + 1% FBS. A volume of 125 μL of the diluted bacterial suspension was added to each well (including controls), resulting in a Multiplicity of Infection (MOI) of 5:1 (bacteria:HT29 cells). The plates were incubated for 60 minutes at 37 °C in a 5% CO_2_ atmosphere. After incubation, the contents of the control wells (without cells) were collected, serially diluted, and plated onto appropriate media to determine the input CFU. In the wells with HT29 cells, the medium was removed, and monolayers were washed three times with 1 mL HBSS to eliminate non-adherent bacteria. To recover adhered bacteria, 100 μL of trypsin was added to each well and incubated for 5 minutes at 37 °C. The resulting cell suspension was recovered using 900 μL of Maximum Recovery Diluent (MRD, Difco, BD), serially diluted, and plated for enumeration. Plates were incubated at 37 °C under appropriate conditions, and viable counts were used to assess each strain’s adhesive capacity. Relative adhesion index (P) was calculated using the formula P = (μ / M) × 100% where P represents a relative log-scale adhesion index rather than a direct CFU-based adhesion percentage; μ is the log-transformed viable count (CFU) of adhered bacteria; and M is the log-transformed viable count (CFU) of the corresponding strain in wells without HT29 cells. To validate the assay’s technical reliability, *L. paracasei* ATCC 334 was used as an internal control. This strain is considered acceptable if its relative adhesion index falls within the validated range of 64–71%, determined through repeated in-house testing. Test has been performed in triplicate (if highly comparable, only one test is shown in the Results section).

### Mouse oral administration, serum GLP-1 measurement, and fasting-refeeding test

All animal experiments were approved by the University of Michigan Institutional Animal Care and Use Committee (IACUC). Two independent cohorts of 20 female BALB/c mice (Jackson Laboratories) were raised until 8 weeks of age before treatment. Within each cohort, mice were divided into two groups of 10 each. The first group served as the control group and was administered *B. subtilis* PY79 spores expressing GFP. The second group was administered PY79 spores expressing Tr 5×GLP-1. The administered strains contained a Cm resistance marker. The spore dosage per mouse was set at 1 × 10^9^ CFU, and the suspension volume was 150 µL per mouse. Spores were directly delivered into the stomach using oral gavage needles. The spore gavage was repeated for three consecutive days (days 1–3). The fecal samples were collected on the third day after gavage treatment and placed in 300 μL of sterile PBS buffer, followed by 15 min incubation, mixture homogenization and centrifugation. The supernatant was spread on LB agar containing chloramphenicol and incubated at 37 °C for 12–16 h to verify colony formation and confirming successful recovery of the spores in the gastrointestinal tract. The first mouse cohort was used for the first serum GLP-1 measurement trial, including fecal bacterial recovery analysis and fasting-refeeding testing, whereas the second mouse cohort was used for the second serum GLP-1 measurement trial.

Serum GLP-1-related peptide measurements were conducted in two trials. In the first trial, blood samples were collected from the orbital sinus on days 3 and 5 following the start of oral administration (day 1). In the second trial, blood samples were collected on days 3 and 7. Approximately 100 µL of blood was collected from the orbital sinus using sterile heparin-free glass capillary tubes and transferred into microcentrifuge tubes. Blood samples were allowed to clot at room temperature for 30 min without addition of DDP-4 inhibitors and were then centrifuged at 3,000 × g for 10 min at 4 °C, after which the serum fraction was carefully collected for downstream analysis. The resulting serum samples were subsequently analyzed by liquid chromatography–mass spectrometry (LC-MS) (Qingdao Saiding Biological Pharmaceutical Co., Ltd., China) for detection of GLP-1-related peptide signals corresponding to the engineering Tr 5×GLP-1 construct.

After confirming recovery of the administered strains, fasting-refeeding tests were performed on days 3 and 5 following the start of oral administration in the first trial. The test was performed where both groups of mice were fasted overnight by removing food at 6:00 PM the day prior (days 2 and 4). At 10:00 AM the following day (days 3 and 5), food was reintroduced. Five mice from each group were randomly selected and blood was collected two hours after feeding via a tail prick method to measure glucose levels with an Accu-Check glucose monitor kit (Roche Diabetes Care GmBh, China). No exogenous glucose was administered during the experiment.

### Evaluation of long-term metabolic effects following oral spore administration

Unlike the short-term studies performed in female BALB/c mice, the chronic oral treatment study was conducted using male diet-induced obese and age-matched lean C57BL/6J mice (Jackson Laboratories) to evaluate the long-term metabolic effects of engineered *B. subtilis* spore administration. A total of 80 mice were used in the study, including 40 obese mice and 40 lean mice. Obese mice were established through high-fat diet feeding (D12492; Research Diets Inc., USA) prior to treatment initiation and exhibited baseline body weights of approximately 30–35 g, whereas age-matched lean mice were maintained on standard chow before treatment initiation and displayed baseline body weights of approximately 24–26 g. During the treatment period, obese mice remained on continuous high-fat diet feeding, while lean mice were switched to the same high-fat diet. All animals were maintained under standardized conditions with free access to water in a temperature-controlled environment (22 ± 2 °C) under a 12 h light/dark cycle. Within both the obese and lean cohorts, animals were randomly assigned into four treatment groups (10 mice per group): blank control (equal-volume water gavage), GFP spore control, Tr 5×GLP-1 spore treatment, and oral semaglutide treatment. Engineered spore groups received daily oral gavage at a dose of 2 × 10^9^ spores per day. The oral semaglutide group received daily oral gavage at 0.345 mg/kg body weight, selected to approximate the clinically used human oral dose of 21 mg/day. The treatment period lasted for 5 weeks. Absolute body weight and body fat percentage were measured weekly throughout the study, and baseline measurements were obtained prior to treatment initiation (week 0). Body fat percentage was measured using dual-energy X-ray absorptiometry (DEXA). Prior to scanning, mice were anesthetized and positioned to minimize movement during image acquisition. Whole-body fat percentage was subsequently quantified from the acquired scans to evaluate treatment-associated metabolic alterations.

### Statistical analysis

All statistical analyses were performed using GraphPad Prism 9 (San Diego, USA). Data are presented as mean ± standard deviation (SD). Statistical significance was evaluated using two-way analysis of variance (ANOVA) or two-way repeated-measures ANOVA as appropriate. For longitudinal measurements, two-way repeated-measures ANOVA was performed using treatment and time as factors, followed by Šídák’s multiple-comparisons test comparing GFP and Tr 5×GLP-1 groups at each timepoint. P value < 0.05 was considered statistically significant.

## Acknowledgments

This work was supported by a sponsor research agreement from Menovo. We would like to express our gratitude to members of the Li lab at the University of Michigan.

## Data Availability Statement

The data supporting the findings of this study are publicly available in Zenodo at https://doi.org/10.5281/zenodo.20187202.

