## Supplementary File 1 for "Gut delivery of pentameric GLP-1 using genetically engineered *Bacillus subtilis*"

### 1. Supplementary Figures

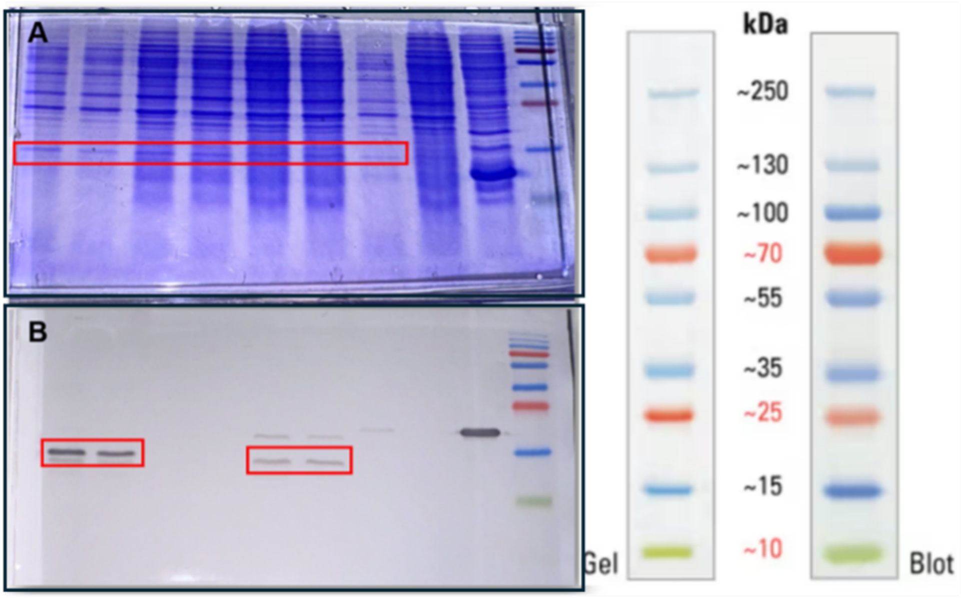


**Figure S1. Expression validation of pentameric GLP-1 constructs.** (**A**) Coomassie-stained SDS-PAGE gel showing expression of pentameric GLP-1 constructs in recombinant *B. subtilis* strains. (**B**) Anti-FLAG Western blot confirming expression of FLAG-tagged pentameric GLP-1 constructs. Boxes in panel **A** and **B** indicate the expected position of pentameric GLP-1 proteins (~15 kDa). Representative data from three independent experiments are shown. **Lane assignment (left to right): 1, 2.** Wt 5×GLP-1 FLAG **3, 4.** Wt 5×GLP-1 **5, 6.** Tr 5×GLP-1 FLAG **7.** Tr 5×GLP-1 (1-7 cultivated with LB containing chloramphenicol) **8.** Control PY79 GFP and **9.** Control PY79.

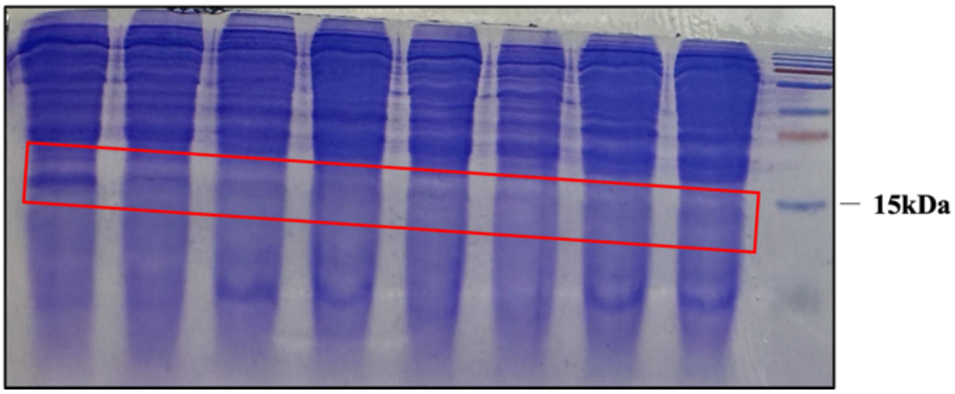


**Figure S2. SDS-PAGE analysis of pentameric GLP-1 secretion before and after chloramphenicol marker excision.** Coomassie-stained SDS-PAGE gel showing TCA-precipitated culture supernatant proteins from recombinant *B. subtilis* strains expressing pentameric GLP-1 constructs. The box indicates the expected migration position of pentameric GLP-1 proteins (~15 kDa). Representative data from three independent experiments are shown. **Lane assignment (left to right): 1.** Wt 5×GLP-1 Flag **2.** Wt 5×GLP-1 **3.** Tr 5×GLP-1 Flag **4.** Tr 5×GLP-1 **5.** Tr 5×GLP-1 (1-5 cultivated with LB containing chloramphenicol) **6-8.** Tr 5×GLP-1 (chloramphenicol marker excised).

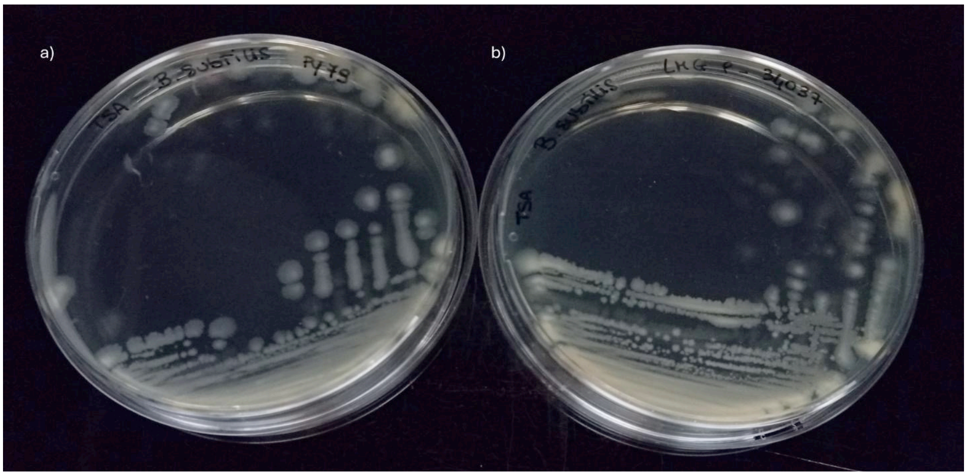


**Figure S3. *B. subtilis* strains isolation and purity check.** The strains **a**) *B. subtilis* PY79 (left), and **b**) *B. subtilis* JH389 (right, in the figure indicated as LMG P-34037), were streaked onto TSA agar plates. Only one morphology was recovered in the plates displaying the typical shape of the *B. subtilis* strain.

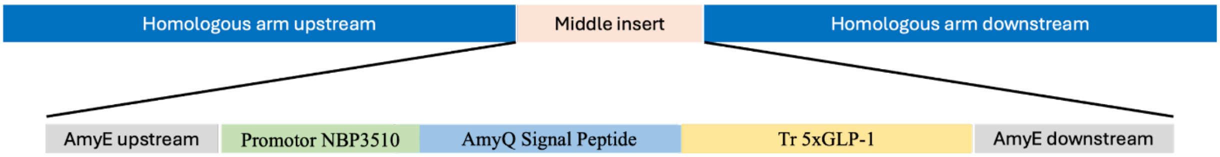


**Figure S4. JH389 Expression Cassette.** The expression cassette in the JH389 strains contains no chloramphenicol resistance marker following the Cre/lox-mediated excision.

#### 2. Supplementary Tables

**Table S1: Primers used in this study**

| Primer | Sequence 5' to 3' | Designation | Description |
|---|---|---|---|
| B685 | TGCTTACGATGTACGACAGGGGG | *amyE*_up_fwd | Sequencing & cPCR |
| B695 | CTACGGCAGCTGATGCTTTTGTAATC | *amyE*_up_rev | cPCR |
| B696 | TACAAAAGCATCAGCTGCCGTAGGATCC | Wt/Tr 5×GLP-1 _fwd | cPCR |
| S170 | TTTGTCATCGTCGTCTTTGTAGTC | Wt/Tr 5×GLP-1_Flag_rev | cPCR |
| B1107 | CGGGGACGTCGACTCTAGACTATTATCTGCCCTTGACGAGCCAAGCG | Wt 5×GLP-1_rev | cPCR |
| B1108 | CGGGGACGTCGACTCTAGACTATTATCTACCATCCACGAGCCAAGCA | Tr 5×GLP-1_rev | cPCR |
| B1104 | TAATAGTCTAGAGTCGACGTCCCCGGG | *amyE*_down_fwd | cPCR |

| | | | |
|---|---|---|---|
| B1101 | GACTACAAAGACGACGATGACAAATAATAGTCTAGAGTCGACGTCCCC | *amyE*_down_fwd | cPCR |
| B690 | CATCCTTGCAGGGTATGTTTCTCTTTG | *amyE*_down_rev | Sequencing & cPCR |
| B741 | GTAATCACTCCTTCTTAATTACAAATTTTTAG | $P_{NBP3510}$_sfGFP_fwd | Sequencing |
| B747 | CTACCGCAGCCAAATATATGAAAAATAGTACATAATGGATTTCCTTACGC | $P_{NBP3510}$_sfGFP_rev | Sequencing |
| S107 | CTGACAGCGTTTCGATCC | $P_{NBP3510}$_sfGFP_rev | Sequencing |
